# Brain connectivity using EEG data

**DOI:** 10.1101/2025.01.26.634935

**Authors:** Anu Aggarwal

## Abstract

The goal of this work was to use EEG data from epilepsy patients and analyze it for connectivity between brain regions using cross-power spectral density (CPSD). We took data from 76 EEG sessions by removing those with epileptic seizures. Then, we calculated CPSD for the 210 electrode pairs for each patient. Initially, we calculated CPSD in the entire sample and then stratified it based on frequency bands related to different brain states. We observed the brain hubs, the default mode network (DMN) and the anti-correlated network attached to the DMN. The connectivity of the brain changed with change in frequency and hence, brain states. EEG analysis with CPSD is a relatively inexpensive, longer duration and more convenient method over fMRI but provides similar information. Similar study can be used to identify brain connectivity patterns while brain is performing a specific task by collecting EEG data for the task.

## I. Introduction

Functional connectivity of the brain is important for understanding brain function. One way to decipher brain connectivity from EEG data is to use cross power spectral density (CPSD) between pairs of electrodes as shown in [2]. Other studies have used fMRI to illustrate the same. fMRI has also been successfully used to detect hubs of information processing in the brain. A limitation of using fMRI is that it is expensive. Another issue is that it cannot provide spectral resolution of the data. Also, it cannot be performed on ambulatory patients. On the other hand, EEG is less expensive, can be performed on ambulatory patients and the data can be analyzed at different frequencies. These frequency bands correspond to different brain states like alpha frequency band with restful wakefulness, theta and delta with sleep and beta and gamma with active information processing. By calculating CPSD, EEG data can help detect connectivity between different brain regions. Similar analysis can be performed in different brain states if we do the spectral resolution of data based on the alpha, beta, theta, delta and gamma frequencies.

With this aim in mind, we calculated CPSD of 10-20 EEG data for a cohort of 129 surface EEG recordings from the temple university hospital (TUH) database [1]. This data was collected as part of ambulatory EEG recording in epilepsy patients over several years. CPSD was used as a measure of connectivity as it indicates not only anatomic but also functional connections between two areas. While analyzing the EEG data, seizure episodes were not considered.

The present work uncovered areas of high connectivity called the hubs as have been identified using fMRI in [5], [6], [8], [34]. It also identified different cortical brain connectivity maps in different brain states corresponding to different EEG wave frequencies. This reinforces the prior findings that the brain connectivity maps change with change in brain functions [33], [36] and even at different times of the day as shown in [18]. The limitation of the study is the limited spatial resolution of the EEG, especially while using 10-20 surface electrodes. Also, the patients were not recorded specifically while performing specific tasks or while sleeping. All the limitations were considered while analyzing the EEG data.

## II. Methods

We analyzed 129 EEGs sampled at 250Hz, by filtering out 60Hz noise and by removing mean and dividing by standard deviation. For further analysis, we removed the EEG recordings with epileptic episodes. For every normal EEG recording, CPSD of each channel with respect to every other channel (210 electrode pairs) was calculated to find the most communicating channels. For CPSD calculation, 4s EEG segments with 50% overlap at a frequency resolution of 1Hz were used. Then, top pairs with respect to their CPSD scores were identified. In the first experiment, we plotted the pairs of channels communicating the most (CPSD > 0.5) for the first 30 frequencies in a single normal patient on a cartoon of the head to easily identify connectivity between different regions. We also color mapped the CPSD for all the 210 electrode pairs (irrespective of CPSD). This uncovered the cortical hubs of information and small world nature of the connections. However, this EEG sample was recorded when the patient was awake.

In the second experiment, we performed a similar analysis on data from all the patients. For this experiment, the 76 EEG recordings were chosen such that the patients were awake, drowsy and asleep at least at some point during the recording and did not experience any seizures. This was done to ensure that we can get enough data for all the brain states but without abnormal activity (seizures). Then, we averaged CPSD for each of the 210 pairs of electrodes from all the 76 patients. This averaged CPSD was color mapped for all 210 pairs of electrodes to visualize their relative connectivity. Of these, the areas with highest CPSD (>0.5) were marked on a cartoon of head for easy visualization. This helped understand connectivity between the brain regions and flow of information.

In the third experiment, we analyzed the 76 normal EEG recordings from the same patients as in the second experiment. In this experiment, we analyzed the CPSD for the 210 pairs of electrodes at different frequencies (alpha, beta, theta and delta frequencies). These frequencies correspond to different brain states (sleep, deep sleep, awake restfulness, awake and active). The resulting CPSD scores for each pair of electrodes at a frequency were averaged across all 76 recordings. The average scores were then plotted as color maps to show the connectivity between the brain regions represented with the electrodes. We also plotted the strongest connections (CPSD>0.5) on a cartoon of the head to visualize the regions connected the most at a particular frequency.

## III Results

Results from the first experiment include analysis of CPSD between all 210 pairs of EEG electrodes for a single patient. Right hand side of Fig. 1 shows a matrix where connection strength between the pairs of channels is shown using colormap. Colorbar in the graph indicates the strength of connection represented by the color. Correlation value of 1 means autocorrelation. Matrix is symmetrical along the diagonal. On the left side of Fig.1, on a cartoon of head with EEG electrodes, the strongest connections (correlation>0.5) are hand drawn lines in green and blue color. Green lines indicate stronger connection between the regions (higher correlation coefficient) than blue lines.

**Fig. 1.**
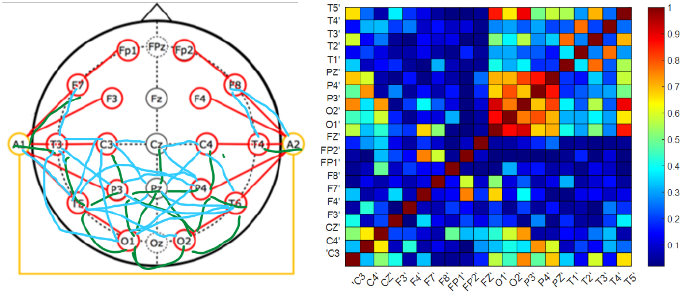
Position of 10-20 electrodes on scalp and correlation between them. In the right figure, nodes with cpsd based correlations more than 50 percent are shown connected with green and blue hand drawn lines (yellow line between A1 and A2 does not indicate correlation). Green lines indicate higher correlation than blue lines. Adjacent nodes communicate more, especially with those in the same hemisphere. Some nodes communicate a lot with neighbors like P3, P4, Pz and Cz. These correspond to cortical hubs identified in earlier studies.

### Small world networks

Most connected regions based on this graph are P3, P4, Pz, C3, C4, Cz, T5 and T6. A notable observation is that most of the connections are between electrodes on the same side of midline. This is consistent with the finding that brain connectivity is a small world network [8]. Remarkably, the frontal cortical electrodes are not communicating much with other electrodes or with each other in this patient. Pz and Cz seem to help in inter-hemispheric communications while frontal region seems to be connected to the central, parietal and temporal through T3-F7 and T4-F8 connections. Rest of the connections had CPSD < 0.5. Hence, these are not shown here.

In the second experiment, we analyzed sEEG data from 76 non-epileptic sessions. All the records were collected over a long period. Therefore, all the patients were in awake, drowsy and asleep states of the brain at least at some point in time. As already mentioned in the methods section, we processed the EEG recordings to remove noise, calculated the CPSD for each electrode with every other electrode. We took the mean of CPSD for all the patients and plotted the graphs in Fig. 2 like in Fig. **1**.

**Fig. 2.**
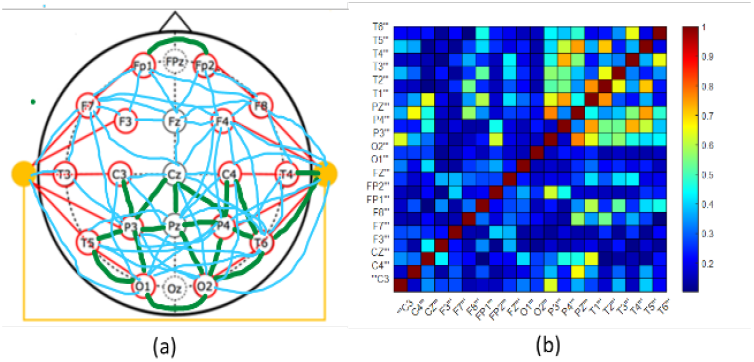
Most communicating pairs (correlation >0.5) of EEG electrodes from a cohort of 76 non-epileptic patients exhibiting awake, drowsy and asleep recording. The second important thing noticed was the top communicating areas in the cortex in a cohort of 76 non-epileptic EEG recording sessions. This cohort was chosen such that all patients are in a mix of awake, drowsy and sleep states. All pairs of electrodes with their correlation of CPSD for this group are shown in the color chart in Fig. 2(b). From this, we plot the connections with correlation >0.5 on a cartoon of the head in Fig. 2(a). There is a similar pattern as seen in data from single patient – more connectivity between areas on the same side of brain, more connectivity with the hubs. Except for that in this cohort, we also see connections between and to frontal electrodes. The reason for this will be clearer when we examine data in frequency bins corresponding to sleep states below.

### Identifying cortical hubs

In these graphs, we can identify the cortical hubs as identified in [5] using the fMRI data. The connections are the strongest on the same side of midline which confirms the brain being a small world network [8]. Here also, the electrodes along the midline, (Cz, Pz and Fz) seem to be helping in inter-hemispheric communication and those along the periphery in anterior-posterior cortical communication.

In addition, there is inter-electrode communication in the frontal region as opposed to little or none in Fig. 1. This could be because in the first experiment, the single patient was in awake state and sedentary. Whereas in the second experiment, the patients are in awake, drowsy and asleep states at least at some point during the record. The latter recordings are over a longer time. Hence, patients might have been more active physically. The results seem to suggest that the frontal lobe is more active as the patient is more active or in asleep state. To explore this further, we analyzed the data by segregating it based on various frequencies.

### Connectivity at different frequencies and brain states

Therefore, in the third experiment, we analyzed the electrodes and hence, the brain areas that communicate the most at a particular frequency range corresponding to the EEG waves – alpha waves (8-12Hz), beta waves (13-30Hz), theta waves (4-7Hz) and delta waves (<4Hz) in the 76 non-epileptic sessions. These EEG wave frequencies are also indicative of the brain state (restful wakefulness, active, sleep, deep sleep respectively) of the patient. After calculating the CPSD for the 210 pairs of electrodes for all 76 recordings, we calculated their average value. The results are presented in Fig. 3 to Fig. 6 – all the channel pair correlation coefficients are shown in color chart on the right and connections between pairs of most connected (correlation>0.5) is shown on cartoon of head on the left. Observations include – insignificant (<0.5) correlation at <1Hz (not shown here), higher correlation between electrode pairs was observed at 1-3Hz, 7Hz, 12Hz than at other frequencies. The channel pairs with significant (>0.5) correlation and hence, information communication to each other in **delta** frequency range were O1-O2, O1-T5, O1-P3, O1-Pz, O2-Pz, O2-T6, Pz-P3, Pz-P4, P3-C3, P3-T5, P4-C4, P4-T6, Pz-Cz, T5-T3, T4-F8, F8-FP2, T3-F7 and F7-Fp1 as shown in Fig. 3. The regions of high connectivity in this figure also include the cortical hubs as seen in Fig. 1 and Fig. 2. In addition, the information flow between frontal and temporal/central electrodes seems to happen along the periphery (T3-F7, T4-F8) more than along the center (Cz-Fz, C3-F3 or C4-F4). And the brain connectivity is like that in a small world network. In the **theta** frequency range, the most communicating pairs of electrodes are O1-O2, O1-T5, O1-Pz, O2-Pz, O2-T6, Pz-P3, P3-T5, P4-C4, P4-T6, Pz-Cz, T4-F8, as seen in Fig. 4. These correlations also span the entire cortex but less so than the delta frequency range. Again, the small-world nature of connections can be seen with stronger connections to nearby nodes, especially on the same side of the midline. And communication between anterior and posterior parts seem to flow along the periphery. Similarly, in the **alpha** frequency range, the strongest correlations are between channels O1-O2, O1-T5, O2-T6, P3-T5, P4-T6 and P4-C4 as seen in Fig. 5. Some weaker correlations are also seen between other cortical hubs and adjoining areas. The cortical areas seem to communicate less in this brain state than in the state related to theta and delta waves. And in the **beta** range, only O1-O2, P3-T5, P4-T6 and C4-P4 are strongly correlated as seen in Fig. 6. This indicates that as the frequency increases, the connectivity between electrodes in the cortical region decreases. This could be because at higher frequencies, the brain regions communicate more with the rest of the body through subcortical regions and spinal cord.

**Fig. 3.**
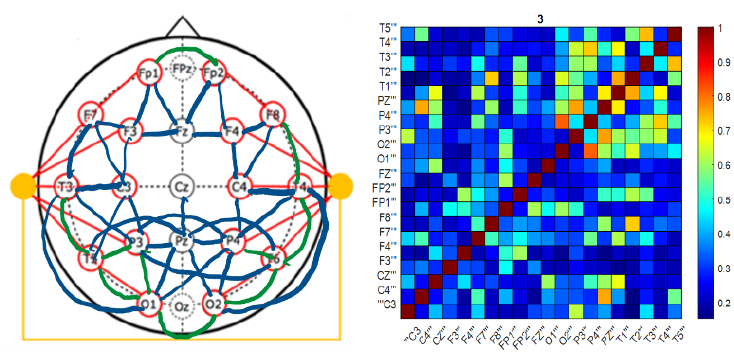
Delta waves (3Hz) – connectivity between different EEG electrodes in 76 non-epileptic EEG epochs. Blue and green lines in the left cartoon only show correlations >0.5.

**Fig. 4.**
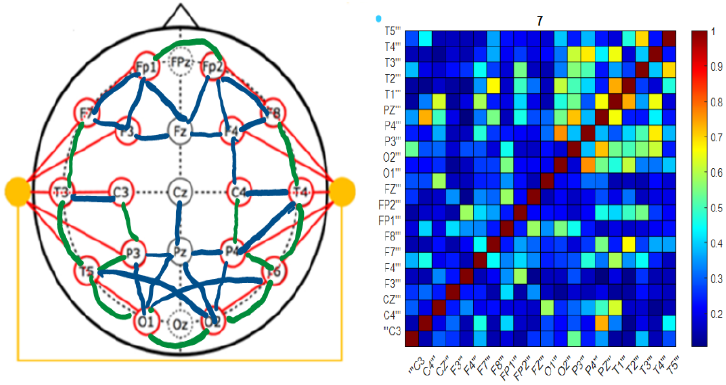
Theta waves (7Hz) - connectivity between different EEG electrodes in 76 non-epileptic EEG epochs. Blue and green lines in the left cartoon only show correlations >0.5.

**Fig. 5.**
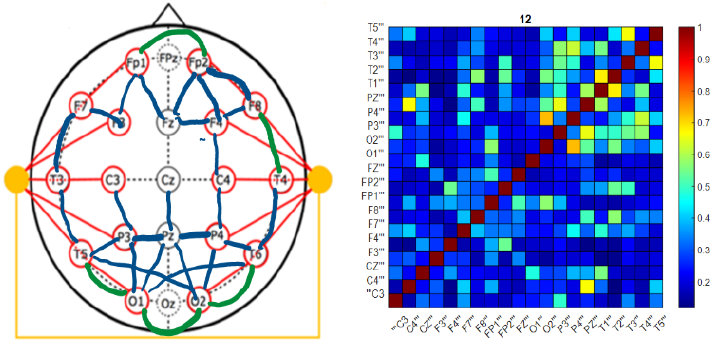
Alpha waves (12Hz) - connectivity between different EEG electrodes in 76 non-epileptic EEG epochs. Blue lines in the left cartoon only show correlations >0.5.

**Fig. 6.**
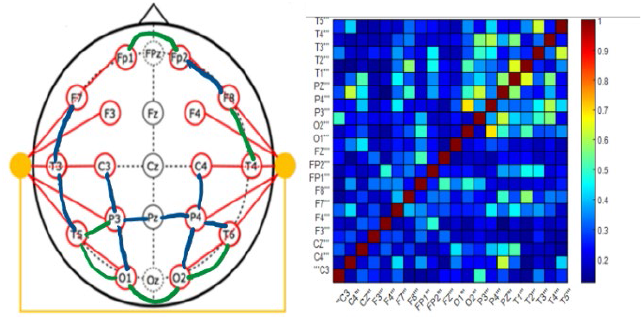
Beta waves (18Hz) - connectivity between different EEG electrodes 76 non-epileptic EEG epochs. Green lines in the left cartoon only show correlations >0.5.

## IV. Discussion

The above results indicate that EEG processing using CPSD analysis is a good way of finding information flow between brain areas. The EEG is a simpler and less expensive method, it can also be used in a mobile setting and for longer duration unlike fMRI or CT. It also highlights the cortical hubs of information in the brain as have already been defined by other imaging methods like fMRI and diffusion tractography. **Cortical Hubs**. According to [5], the structural core of the brain includes the regions - the posterior cingulate cortex, the precuneus, the cuneus, the paracentral lobule, the isthmus of the cingulate, the banks of the superior temporal sulcus, and the inferior and superior parietal cortex, all of them in both hemispheres. Most connected regions based on the graphs in Fig. 1-Fig. 6 seem to be regions P3, P4, Pz, C3, C4, Cz, T5 and T6. These all correspond to the 8 cortical hubs – Cuneus (BA^1^19), precuneus (BA 17, 18), paracentral lobule (BA6), superior parietal cortex (BA7), inferior parietal cortex (BA39, 40), posterior cingulate cortex (BA23), isthmus of the cingulate (BA31) and banks of the superior temporal sulcus (BA22) identified in [5] using MRI and Diffusion tractography data. According to [3] and [4], P4 and Pz electrodes correspond to BA19 which corresponds to the cuneus. C3 and C4 correspond to BA 6 and 7 which correspond to paracentral lobule and superior parietal cortex. Other cortical hubs include BA 39 and 40 which are recorded by P3 on surface EEG. The superior temporal sulcus (BA22) corresponds to T5 and T6 electrodes in surface EEG. The posterior cingulate cortex (BA23), and isthmus of the cingulate (BA31) correspond to Cz electrode. This implies the calculation of CPSD of EEG can correctly identify the cortical hubs in brain. This is summarized in Table 1 below.

**Table 1.** Brodmann areas and electrodes associated with cortical hubs.

| S.No. | Cortical Hub | Brodmann area | Electrode of sEEG |
| --- | --- | --- | --- |
| 1 | Posterior cingulate cortex | 23 | Cz |
| 2 | Precuneus | 17, 18 | Pz |
| 3 | Cuneus | 19 | P4, |
| 4 | Paracentral Lobule | 6 | C3, C4 |
| 5 | Isthmus of the cingulate | 31 | Cz |
| 6 | Banks of the Superior Temporal Sulcus | 22 | T5, T6 |
| 7 | Inferior Parietal Cortex | 39, 40 | P3 |
| 8 | Superior Parietal Cortex | 7 | C3, C4 |

### Memories, connectivity, and information processing

From the strongest connections between brain areas, it seems that information flows from sensori-motor areas and visual cortex to the visual sketch pad. From visual areas, it also spreads to attention area, meaning construction area and long-term memory. Visual sketch pads are in turn connected to long-term memory and attention area. Basal ganglia are connected to the attention area, visual sketch pad and meaning construction area. This is illustrated in detail in Table 2. Primary and secondary visual areas on both left and right hemispheres seem to be correlated due to simultaneous activation of the two from external visual stimuli (O1-O2). This means visual sketch pad gets visual input and other sensory inputs, processes the information, and provides it to long-term memory and attention area. In long-term memory, it could form spatial information for episodic memory. In the attention area, it can be important to draw attention to important areas perceived by sense organs. Meaning is constructed based on visual input and giving attention to the area. Basal ganglia guide motor actions based on direct visual inputs and those routed through the attention area and meaning construction area.

**Table 2.** Showing region pairs having highest cpsd correlation in our data of 129 patients. **Col2 and 3 show the anatomical brain area associated with the electrodes and the function of the area.**

| EEG Electrode pair | Function and anatomical area related to first electrode in the pair | Function and anatomical area related to second electrode in the pair |
| --- | --- | --- |
| P3-Pz | Acalculia - BA 39 Angular Gyrus - Meaning Construction | Agnosia, Apraxia - BA 7 Attentional Shifting Area - Perseverance |
| P4-Pz | visual spatial sketch pad (7) | Agnosia, Apraxia - BA 7 Attentional Shifting Area - Perseverance |
| O2-P4 | V1, V2 (17, 18) Rt | visual spatial sketch pad (7) |
| Cz-Pz | Basal Ganglia (Thalamic Efferent VA) Substantia Nigra (Thalamic Efferent VL) | Agnosia, Apraxia - BA 7 Attentional Shifting Area - Perseverance |
| P4-T6 | visual spatial sketch pad (7) | Long-Term Memory (21), Facial Recognition - Emotional Content |
| O2-T6 | V1, V2 (17, 18) Rt | Long-Term Memory (21), Facial Recognition - Emotional Content |
| O2-Pz | V1, V2 (17, 18) Rt | Agnosia, Apraxia - BA 7 Attentional Shifting Area - Perseverance |
| O1-O2 | V1, V2 (17, 18) Lt | V1, V2 (17, 18) Rt |
| Cz-P4 | Basal Ganglia (Thalamic Efferent VA) Substantia Nigra (Thalamic Efferent VL) | visual spatial sketch pad (7) |
| Cz-P3 | Basal Ganglia (Thalamic Efferent VA) Substantia Nigra (Thalamic Efferent VL) | Acalculia - BA 39 Angular Gyrus - Meaning Construction |
| O1-P3 | V1, V2 (17, 18) Lt | Acalculia - BA 39 Angular Gyrus - Meaning Construction |
| FP1-FP2 | Verbal, Episodic Retrieval Visual Working Memory (10) | Face & Object Processing - Emotional Inhibition, Verbal Episodic Memory (supra-orbital region) |
| C4-P4 | sensorimotor area, short term memory (3, 4, 40) | visual spatial sketch pad (7) |

<sup>1</sup> Brodmann area

**Frequency-based analysis** of the data indicates that there is much higher correlation between brain areas and hence, higher information flow from one area to another at delta frequencies. This is followed by the theta frequencies. This gradually reduces at alpha frequency range. Eventually, in beta frequency range, there is very little correlation between brain areas. This indicates that most of the inter-regional communication in the cortex happens during NREM sleep, especially during delta wave or slow wave sleep and theta wave or stage I sleep. While at higher frequencies, the brain communicates more with periphery as the body is actively involved in sensori-motor tasks.

In the delta frequency range, there is a network formed by connections between F4, F3, Fz, C3, C4, P4, P3, Pz, T3 and T4 as seen in Fig. 3. This corresponds to the Default mode network [20] (DMN) that is considered a task negative network, showing increased metabolic demand during the ‘baseline’ state of the brain. It is active when individuals are not focused on the external environment but involved in intrinsic activity such as autobiographical memory and future envisioning. It includes cortical areas like PCC, RspC, inferior parietal lobule (IPL), medial prefrontal cortex (mPFC), parts of the hippocampal formation and the temporal lobe. This is not observed in the beta frequency range (Fig. 6). Even in alpha frequency range (Fig. 5), this is disrupted.

It is coupled to an anticorrelated network (ACN) that is involved in goal directed task performance. The latter includes areas like the intraparietal sulcus, frontal eye fields, middle temporal area, supplementary motor area and insula (BA 40, 21, 8, 6, 13). This corresponds to EEG electrodes C3, C4, T3, T4, F3, F4, Fz and Cz. Subregions of this ACN overlap with the dorsal attention network (intraparietal sulcus, frontal eye fields, and supplementary motor area) and with the ventral attention network (insula). The anterior and posterior midline node of the DMN is linked to these ACNs. These are linked to the DMN in different graphs. These are more consistent across brain states seen in Figs. 3-6. This implies, the ACNs are active across brain states but DMN will be more active in the resting states.

The results also indicate that the cortical brain areas communicate a lot more at lower frequencies that correspond to sleep states. This could mean that consolidation of information will happen more in asleep than awake states as the brain cortical areas are communicating more. And that in awake states, most of the body movements or spinal control is exerted through the sensori-motor areas and the basal ganglia with little involvement of frontal cortex (related to planning). Similar observations have also been made in modelling with human and primate data. In [31], authors observed that there is reduction in inter-regional correlations and variability when the subject is performing a task.

### Frequency and fidelity of long distance communication

Thus, the brain seems to use a higher frequency for longer distance communication – perhaps because it can send more information within a short time duration. While it reserves the lower frequencies for information consolidation and shorter distance communication as within the cortex. From our knowledge of communication systems, we know that at higher frequencies, the attenuation of information is higher than that at lower frequencies. Therefore, in communication systems, we use higher frequencies for shorter distances and lower frequencies for longer distances. While lower frequency waves may be blocked or weakened by obstacles, high frequency waves can often pass through them with minimal attenuation, so the communication science uses the higher frequencies for sending messages in urban areas where there are many obstacles on the way. Generally, low-frequency transmissions can travel greater distances before losing their integrity and they can pass through dense objects more easily, but less data can be transmitted over these radio waves. This could indicate that the memories are formed at higher fidelity but slowly, hence, need a longer time to consolidate. While the bodily movements and sensory perception require quick processing even if the fidelity of the signal is low (maybe there are other mechanisms for applying corrections to the signal).

In [7], it is mentioned that delta frequency sets the functional bias in large neural networks in brain while moderate frequency alpha helps in transferring around in brain while still higher frequency beta and gamma frequencies transfer to longer distances. And that beta is associated with motor tasks. As mentioned in pg 1103 of [7], gamma oscillations do not seem to increase information processing by different cortical areas due to synchronization. But the stimulus perception may be related to gamma because this study does not investigate stimulus perception. The stimulus perception will increase inputs from periphery to sensory areas of the brain without changing interconnection between different sensory areas. But in [7], they used the 20-60Hz range for gamma frequency. The gamma band activity increases with presentation of familiar sounds but not novel sounds as it is involved in top-down attentional processing. Gamma activity has also been involved in encoding and retrieving from memory in hippocampus and entorhinal cortex by formation of a brain workspace. This was not observed in our study as we are not observing gamma frequency band and only observing interconnections between cortical areas. We are not observing the cortico-subcortical connectivity.

### Thalamus as a switch

Another interesting observation from [18] is that there is diurnal variation in cortical connectivity. They demonstrated higher interhemispheric parietal connections in the morning than afternoon which they thought could be due to planning in the morning for the entire day. They also observed higher cortical connectivity between frontal and parietal regions during afternoon which they surmise to be due to the two regions interacting to generate and monitor a vivid experience of mental imagery. Overall, they found it to be related to retrospective introspection to explore the relationship between the ongoing subjective experience of participants and the key characteristics of their brain during resting state. In [20], there has been reduced bilateral para hippocampal gyrus and right inferior temporal gyrus connectivity to default mode network (DMN) during S1 to level of no contribution during slow wave sleep. Whereas, connectivity of medial PFC, and bilateral posterior cingulate cortex (PCC) and retro splenial cortex (RspC) to the DMN only decreases from wakefulness to slow wave sleep. But in this study, they only observed default mode network areas. They also used cross correlation analysis and independent component analysis of fMRI and EEG data. Similar findings regarding dissociation of medial pre-frontal cortex from the DMN during deep sleep was also observed in [21] which interpreted it as conscious mentation requires DMN network to be intact. This is the opposite of what we observed in this study. This has been seen to be associated with thalamus acting as a switch and regulating the flow of information from periphery differentially depending upon the brain state in [37].

### Small world networks

The data analysis in our study as shown above also demonstrates higher connectivity between areas placed close to each other, especially those in the same cerebral hemisphere. This is in line with small world nature of brain connectivity as shown in [5].

### Changes in brain connectivity

There are some more notable observations from other studies of brain connectivity. According to [17], connectivity between parts of fronto-temporal cortex is increased at theta range and decreased in alpha range in patients with autism spectrum disorder. Whereas [19] proposes that there is increased connectivity between subcortical nuclei like thalamus and caudate nucleus and the sensory cortex in patients with autism spectrum disorder as compared to matched controls. According to [10], the brain exhibits repeating nonrandom patterns of activities that correlate with, and strongly influence, behavior. With a specific study of hand movements and eye movements [16] influencing the cortical connectivity in cerebral and cerebellar areas involved in the movements like the sensorimotor and visual areas. Understanding the states of the brain, their mechanisms of generation, and how they interact with sensorimotor and cognitive processing is essential to achieving a thorough understanding of neural mechanisms of natural behavior [11]. This study can help develop models to predict those changes.

Change in brain connectivity related to brain state might be due to change in long range connections between regions[12]. The studies of brain area connectivity have also been used to predict individual potential at learning new things like motor tasks as in [25]. Or in understanding the state of delirium [26] where there is a decrease in the anticorrelation between the default mode network and task positive regions, excessive internal connections in the posterior default mode network and a complex imbalance of internal connectivity in the anterior default mode network. There is a loss of reciprocity between the default mode network and central executive network associated with defective function in the salience network which can lead to aberrant subcortical neurotransmission related connectivity and striato-cortical connectivity. According to [23] and [24], same connection network may be part of different functional dynamics and hence, play role in adaptation meaning modify connections with the function for which the network is being used. Similarly, [27] also proposes that there are different brain states of functional connectivity that manifest different states of consciousness. Studies like [32] suggest that there is a single functional connectivity state, the latent state that manifests in different connectivity states during different mental tasks and that this might also be linked to the intelligence of the subject.

## V. Conclusion and future work

In this paper, we used the CPSD method to analyze EEG data to identify the cortical hubs with high connectivity in brain. These results are like those obtained using more complicated, expensive, and restrictive methods like fMRI and CT. We also identified electrodes that communicate more at frequency bands corresponding to EEG waves. It was observed that there is higher connectivity between areas on the same side of the brain. Most of the activity during sleep was seen in cortical areas involved in the formation of short term and long-term memories. In contrast, there is very little transfer of information between cortical areas during restful wakefulness (alpha frequency range) and active brain function (beta frequency range).

There are several applications of this work.

1. Studies like ours can be useful to determine AD, MCI where one of the causes of reduced cognitive ability is network disruption, especially at the hubs. As in these diseases, functional connectivity between areas is disrupted [9], [35]. It can also be useful in determining changes in functional connectivity of brain areas leading to audio-visual hallucinations in diseases like Schizophrenia [14]. In [15], the authors show the value of studying cortical connectivity to identify mental fatigue while performing a task over a period. These can guide the development of models that can be used to test treatments or interventions in patients.
2. We can also identify patterns of connectivity in the brain during a task. Thus, can find signature EEG patterns during a task as has been done in [13], [23]. Based on which we can identify what a person is doing or intending to do based on their EEG alone. This will require EEG data from people doing specific tasks. For instance, in [22], authors have shown impact of negative and positive emotional states on neuronal firing in prefrontal cortex, temporal and parietal cortex and orbitofrontal cortex and tried to infer their role in emotional processing.
3. Like in [28], we can build models of the brain states and understand their transitions under constraints or to test drugs that might be useful in implementing these transitions and for various other medical purposes.

Limitations. One of the limitations was that the data was taken from the Temple University Hospital EEG cluster where it was collected from epileptic patients either during routine clinical examination or while carrying on with their routine work and not from patients performing specific tasks or in specific brain states. Another limitation was that there were no recordings from the subcortical structures like the thalamus and the basal ganglia. If we could collect EEG data from these as well, then CPSD based analysis might help in identifying the connections and impact on them in ailments like stroke. This was observed in [29], which demonstrates how the clinical course of patients with stroke can be identified by examining changes in brain connectivity maps. Another limitation of the EEG is its limited spatial resolution. Because of which we could not identify 180 separate areas in the cortex as identified in [5]. However, the EEG does provide better opportunity for spectral analysis which is related to brain states and was used in the current study. This could be improved by co-registering fMRI and EEG data. The EEG data for this study was from epileptic patients. Even though the epileptic episodes were excluded from the analysis, there is still a possibility that the brain networks could be altered due to the underlying morbidity.

## Footnotes

1 Brodmann area

## Notes

### Competing Interest Statement

The authors have declared no competing interest.

